# Phylogenetic detection of protein sites associated with continuous traits

**DOI:** 10.1101/2025.07.22.665918

**Authors:** Louis Duchemin, Gerard Muntane Medina, Bastien Boussau, Philippe Veber

## Abstract

Comparative genomic data can be used to look for substitutions in coding sequences that are associated with the variation of a particular phenotypic trait. A few statistical methods have been proposed to do so for phenotypes represented by discrete values. For continuous traits, no such statistical approach has been proposed, and researchers have resorted to sensible but uncharacterized criteria. Here, we investigate a phylogenetic model for coding sequences where amino acid preferences at a site are given by a continuous function of a quantitative trait. This function is inferred from the amino acids and the trait values in extant species and requires inferred point estimates of ancestral values of the trait at internal nodes. For detecting sites whose evolution is associated with this trait, we use a significance test against the hypothesis that amino acid preference does not depend on the trait.

This procedure is compared to simpler strategies on simulated alignments. It displays an increased recall for low false positive rates, which is of special importance for performing whole-genome scans. This comes however at a much higher computational cost, and we suggest using a simple test to filter promising candidate sites. We then revisit a dataset of alignments for 62 species of mammals, using longevity as a phenotypic trait. We apply our method to three protein families that have previously been proposed to display sites associated with variation in lifespan in mammals. Using a graphical representation extracted from the detailed phylogenetic analysis of candidate sites, we suggest that the evidence for this in the sequence data alone is weak.

The proposed method has been added to our Pelican software. It is available at https://gitlab.in2p3.fr/phoogle/pelican and can now be used with both discrete and continuous phenotypes to search for sites associated with phenotypic variation, on data sets with thousands of alignments.

## 1 Introduction

Living organisms show a fascinating diversity in their shapes, lifestyles, life history traits, and behaviours. They have acquired these varied phenotypes during billions of years of evolution. Nowadays, comparative genome data offer the possibility of uncovering genotypic changes associated with phenotypic changes. This opportunity has motivated a lot of methodological development and empirical studies. Several methods have been developed to study rates of sequence evolution in coding and non-coding sequences (e.g., [Lartillot and Poujol, 2011, Kowalczyk et al., 2019, Wu et al., 2017, Gemmell et al., 2024, Goldman and Yang, 1994, Yang, 2007]). These approaches assume that concomitantly with phenotypic changes, some areas of the genome that were under selective constraints could become less constrained, and some other areas could follow the opposite trend. To find genotype-phenotype associations, they infer rates of sequence evolution and compare them to rates of phenotypic evolution [Lartillot and Poujol, 2011] or to phenotypes annotated along a phylogeny [Kowalczyk et al., 2019, Wu et al., 2017, Gemmell et al., 2024, Goldman and Yang, 1994, Yang, 2007]. In particular, *d_N_ /d_S_* approaches [Goldman and Yang, 1994, Yang, 2007] have been widely used to find protein-coding genes or sites associated with changes of a discrete trait along a tree. Apart from approaches based on rates, another type of approach, the *profile* methods, looks for an association between preferred amino acids and phenotypic states (e.g., [Tamuri et al., 2009b, Parto and Lartillot, 2018, Duchemin et al., 2022]). These methods model amino acid preference at a site and given a particular phenotypic value as a profile (Fig. 1), i.e., parameters describing the fitness contribution of having particular amino acids at that site. The consideration of amino acid profiles leads to distinguishing the direction of selection from its intensity. Changes in the direction of selection correspond to changes in the relative fitness of different amino acids; for instance, profiles A and B in Fig. 1 differ notably in that the red amino acid has a higher fitness than the yellow amino acid in profile B but not in profile A. Changes in the intensity of selection at a site make fitness differences between amino acids larger or smaller, but do not change the relative order of the fitnesses. In this manuscript, we focus on changes in the direction of selection associated with phenotypic changes, i.e., we are interested in changes in profile space associated with phenotypic changes, and do not consider changes in selection intensity. We acknowledge that changes in the intensity of selection may also be associated with phenotypic changes, but leave it to future work to detect such effects. Of note, *d_N_ /d_S_* approaches [Goldman and Yang, 1994, Yang, 2007] do not distinguish between the intensity and the direction of selection because they do not rely on amino acid profiles. They can thus be used to detect both types of changes.

**Figure 1:**
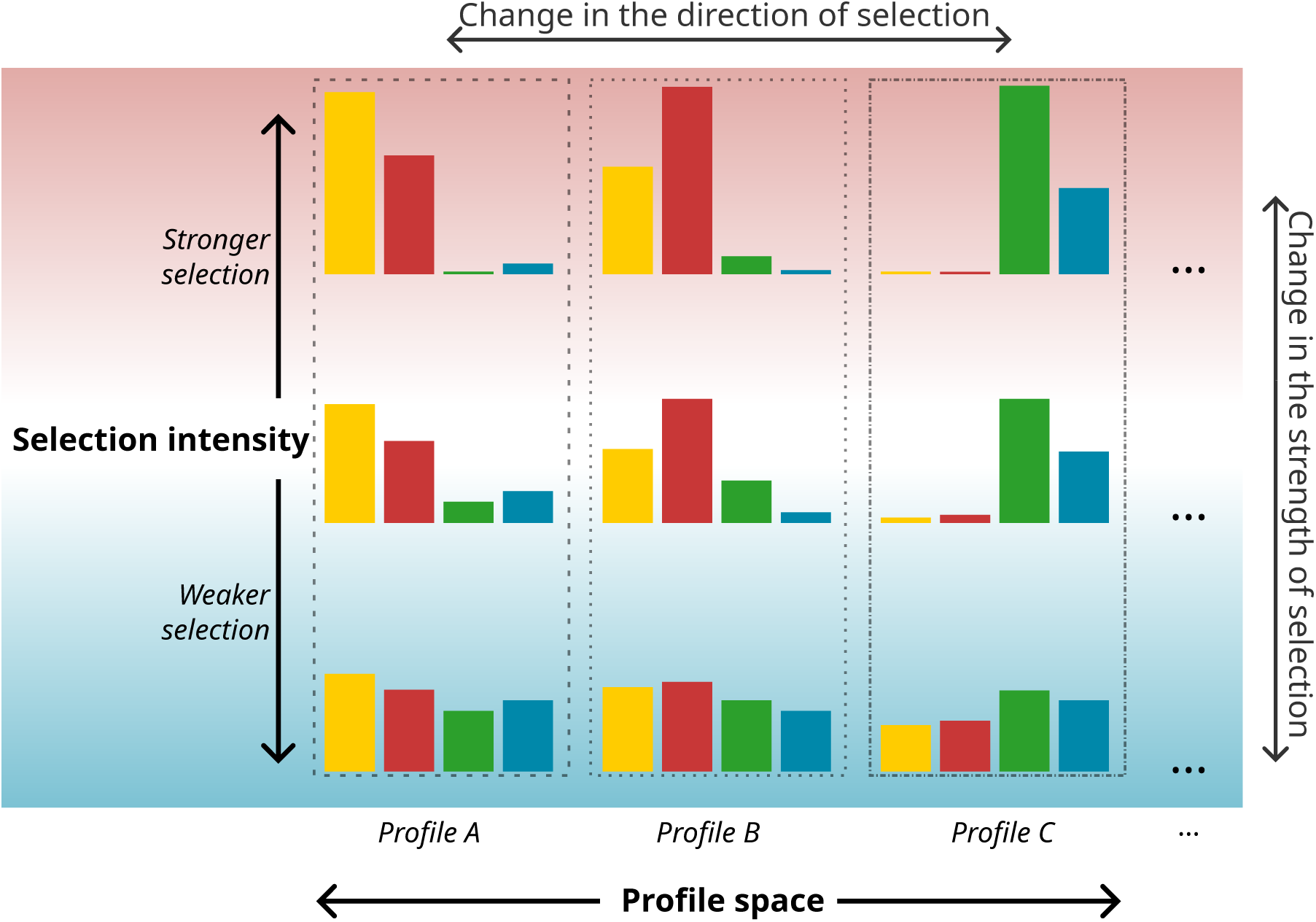
Amino acid fitness profiles describe the direction and strength of selection at a protein site. 3 amino acid profiles are represented, with bars in 4 colors representing fitness values for an alphabet with only 4 different amino acids. 3 levels of strength of selection are represented. Under weaker or stronger selection, amino acid profiles tend to be flatter or more heterogeneous respectively, but the relative orders of the amino acid fitnesses is conserved.

We have shown that some profile methods, in particular the method Pelican which re-implements a model by [Tamuri et al., 2009b], could be as powerful and faster than *d_N_ /d_S_* methods [Duchemin et al., 2022] to detect profile changes associated with changes in discrete phenotypes. Of note, all of these methods, including the new approaches we present in this paper, do not attempt to identify causal relationships between phenotypic traits and genotype: they merely identify correlations between them, which can be interpreted in different ways.

So far profile methods have been used with discrete annotations of a phylogeny, whereby each branch is linked to a particular phenotypic state. Such annotations are ill-suited for continuous phenotypes, because they require discretizing a continuous distribution into categories. There are several ways to do so: for instance, to discretize into two categories, one could split at the mean or the median, or any other quantile. A priori, there is no reason to pick one discretization method rather than another. To circumvent this difficulty, some researchers have relied on two steps. First, putative sites where particular amino acids are associated with phenotypic values are identified by focusing on species with the most extreme phenotypic values. Second, species with less extreme phenotypic values are used to validate the sites identified in the first step. This pioneering approach has been used to identify sites and genes associated with increased longevity in primates and mammals [Muntané et al., 2018, Farré et al., 2021], but it has some drawbacks. First, it still depends on arbitrary thresholds to identify “extreme” phenotypic values. Second, it can only detect sites where all taxa at the high end (respectively, low end) of the phenotypic value have the exact same amino acid.

In this report, we introduce two new models that are generalizations of the discrete model from [Tamuri et al., 2009a], to look for sites associated with continuous phenotypes in the Pelican software package. Model *C_S_* links a continuous trait with amino acid preference at a site using a sigmoid function; Model *C_L_* does so using a multinomial logistic regression. We compare their performance against the [Farré et al., 2021] method and against the original discrete variant (model *D*) where trait values are discretized in different ways, using simulations to generate protein sequence alignments under a mutation-selection model. Simulations are run along the 62-species mammalian phylogeny used in [Farré et al., 2021], using longevity measures as an example for a continuous trait. We show that the continuous variants perform better than the other methods, while exempting from the need to determine a discretization procedure for the trait values. Model *C_L_* performs better than Model *C_S_*, but all models would need a larger number of sequences to reach a false positive rate compatible with genomic screens. We then apply model *C_L_* to three protein alignments with 61 sequences that had been proposed to contain sites associated with variation in lifespan in mammals [Farré et al., 2021]. We discuss the results obtained in relation to the findings in [Farré et al., 2021], and propose a graphical representation based on our model parameters and reconstructions of the detailed history of each candidate site to better assess their relevance.

## 2 Results

### Detecting sites whose evolution is associated to a continuous phenotype

We adapted a method initially introduced by [Tamuri et al., 2009a] to detect shifts in amino acid preferences at a protein site associated with past variations of a discrete covariable (typically a phenotypic trait). The method assumes that we observe the residue at the protein site for a set of extant species, and that the phylogenetic tree and ancestral values of the covariable are known. It uses a continuous-time Markov chain (CTMC) to model the evolution of a site along the branches of a phylogenetic tree, and considers two variants of this model. In the first variant, called homogeneous model, all branches share the same rate matrix for substitutions (i.e., the same infinitesimal generator of the CTMC); in the second variant, called heterogeneous model, each branch is associated with a discrete value, and branches with different values may be associated with different substitution rate matrices. Both models have parameters that need to be inferred from the observed amino acids in extant species, and this is achieved by maximum likelihood. [Tamuri et al., 2009a] uses a General Time Reversible (GTR) parametrization of substitution matrices, where rates are decomposed into so-called exchangeabilities and equilibrium frequencies. Exchangeabilities are assumed to be known (they are taken from the WAG matrix [Whelan and Goldman, 2001]) so estimating rates boils down to estimating a vector of 20 amino acid frequencies expressing the preferred amino acids at the considered site. Finally, a significance test is performed to test against the null hypothesis stating that substitution rates are independent from the phenotypic trait values. We adopt a likelihood ratio approach where the likelihood under the homogeneous model is compared to that of the heterogeneous model where equilibrium frequency vectors may vary between phenotypes. Similar to [Tamuri et al., 2009a] we compute a p-value by assuming the distribution of the log ratio statistic approximately follows a *χ*^2^ distribution. The procedure is repeated on each site of a protein independently.

We previously reimplemented the model by [Tamuri et al., 2009a] in our program Pelican and benchmarked its performance against alternative methods [Duchemin et al., 2022]. Our contribution in this report is to consider the case where branches are annotated with continuous phenotypic values instead of discrete values. We provide two models that relate a continuous trait to amino acid frequencies, and present them in details in section 4.4. Model *C_S_* is a smooth interpolation between two asymptotes corresponding to amino acid frequencies estimated for extreme values of the phenotypic trait. It is expressed as a sigmoid vector function *Y_S_* of the trait value *t*:

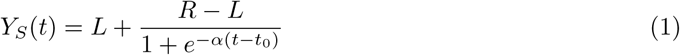

where *L* and *R* are left and right asymptote respectively, *t*_0_ is the trait value at the inflection point of the sigmoid and *α* a slope. Model *C_L_* is based on the combination of a linear function and a softmax link function, as in multinomial regression models :

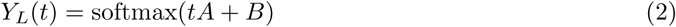

where *A* and *B* are two real vectors of size 20. The softmax function maps any real valued vector *x* to a vector of probabilities that sum to 1. The *i*^th^ coordinate of this vector is defined by:

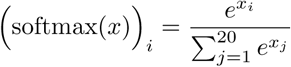

Figure 2 illustrates how the two formulations fit on a simulated example. While they can be made relatively close to each other on the interval spanned by trait values (red rectangle in the figure), their asymptotic behaviour differs substantially: model *C_S_* converges to its asymptotes for extreme values of the trait, whereas model *C_L_* converges to Dirac distributions where only one amino acid is allowed. As a result, the frequency of some amino acids may have a non monotonous dependence to the trait value. This is for instance observed for the brown amino acid, whose frequency first goes up as the trait value increases, then goes down. The motivation behind model *C_L_* is that it can be seen as an approximation of *C_S_* while using fewer parameters than model *C_S_*. The lesser number of parameters may provide added statistical power when performing the likelihood ratio test, as a trade-off for a reduced model fit.

**Figure 2:**
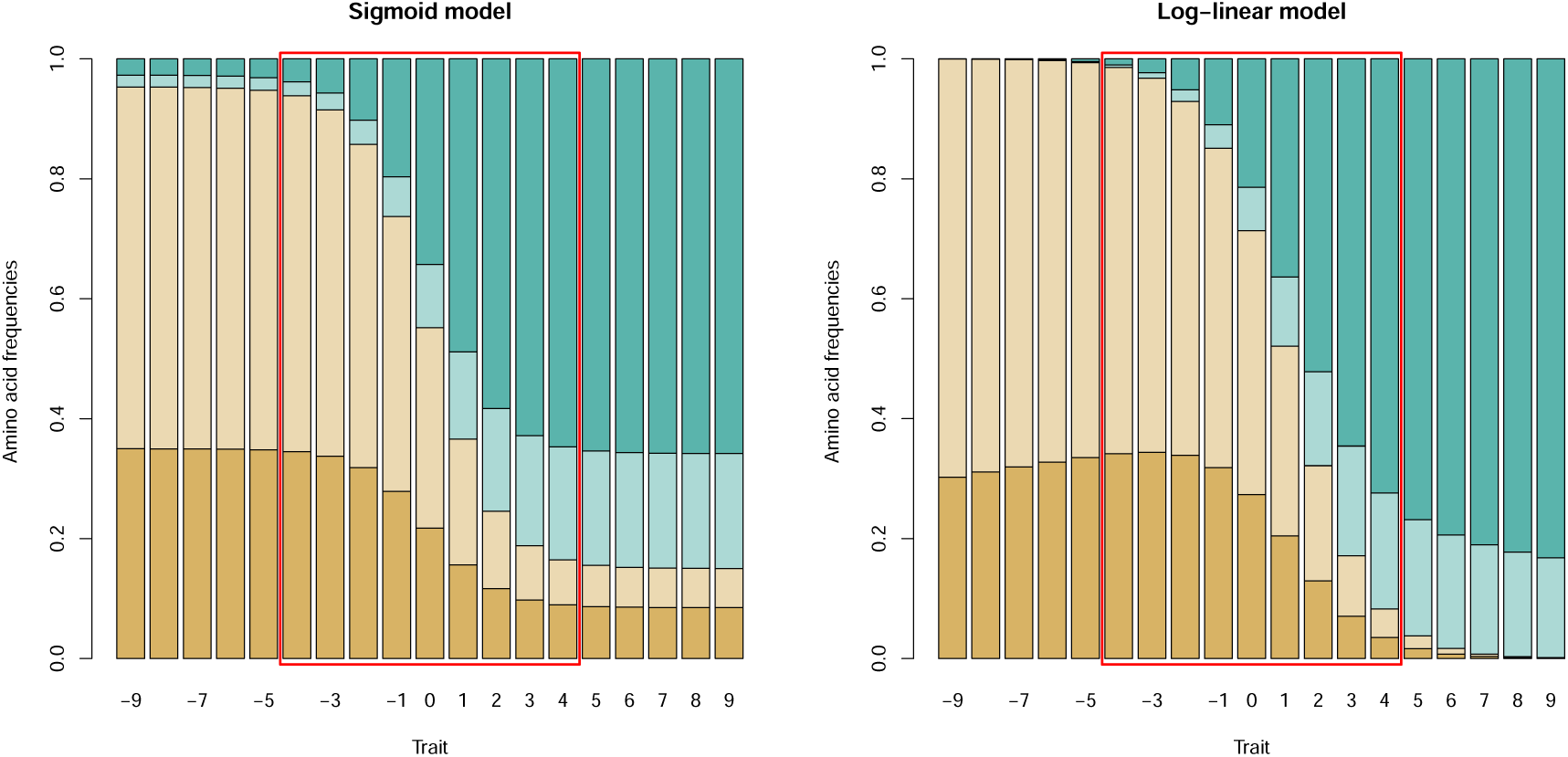
Interpolation of amino acid frequency profiles along a continuous trait using sigmoid or log linear functions. These graphs illustrate the kind of dependency that can be expressed between trait values and amino acid frequency vectors, for each model. They were obtained by simulating a trait-dependent site with 4 amino acids under the sigmoid model, and then fitting each model on the amino acids observed at the leaves. The red rectangles show the interval spanned by trait values; frequency vectors outside of this interval show how each model extrapolates the relation between trait and amino acid frequencies.

### Validation and calibration study

We validated our implementation by simulating alignments under model *C_S_* and used them for two purposes. First, we ensured that when using trees with an increasing number of leaves, the estimates for the various parameters of *C_S_* converged towards their true value. The results are shown in supplementary figure 1 and suggest that the sigmoid slope and asymptote amino acid frequency vectors require 200 to 500 observed species to be robustly estimated from the data. This is a high but not infeasible requirement at present, but in any case it should be stressed that rejecting a null hypothesis may not require a precise estimate of parameters. Second, we evaluated the calibration of the likelihood ratio test under the null hypothesis. We performed simulations under the homogeneous model and along a particular tree (introduced below) with 62 species. For added realism, we used the mutation-selection framework to simulate sites, using empirically determined fitness profiles for amino acids, as implemented in [Duchemin et al., 2022]. Evolutionary rates were tuned to match the amino acid diversity observed in the columns of the empirical alignments used in [Farré et al., 2021] (see supplementary figures 2 and 3). We report in supplementary figure 4 the distribution of calculated *p*-values for model *C_S_* and model *C_L_*. In both cases, we observe that *p*-values are reasonably calibrated, given the size of the sample and despite simulating under a codon mutation-selection model, but inferring under an amino acid model. The peak at 1 for model *C_S_* is caused by instances where Pelican fails to optimize the likelihood of the heterogeneous model. Model *C_L_* displays a small bias towards small values, which means that significance values as well as false discovery rates that can be estimated from them using e.g. the Benjamini Hochberg procedure might be too optimistic. This observation is important to consider when interpreting empirical data analyses.

### Continuous models display improved performance on a simulated benchmark

To evaluate the merits of the different approaches to detect sites associated with continuous traits, we implemented another simulation where we generated sequence alignments with two types of sites: negative (*H*_0_) sites for which the model of sequence evolution is homogeneous across the phylogeny, and positive (*H_A_*) sites for which the fitness of an amino acid in the mutsel model is a sigmoid function of a continuous trait evolving along the phylogeny. We next assessed the performance of several methods by reporting their ability to detect positive sites (recall) under a range of false positive rate thresholds. It is essential here to control false positive rates as those methods are typically meant to be used for genome-wide scans involving millions of site-wise tests.

The simulations were sized according to a study that sought convergent substitutions in relation to longevity in a phylogeny of 62 mammalian species [Farré et al., 2021]. We used the *longevity quotients*, i.e. the log of the actual longevity of a species and the longevity predicted assuming an allometric scaling law between longevity and body mass. The phylogenetic tree and the longevity trait are represented in figure 3.

**Figure 3:**
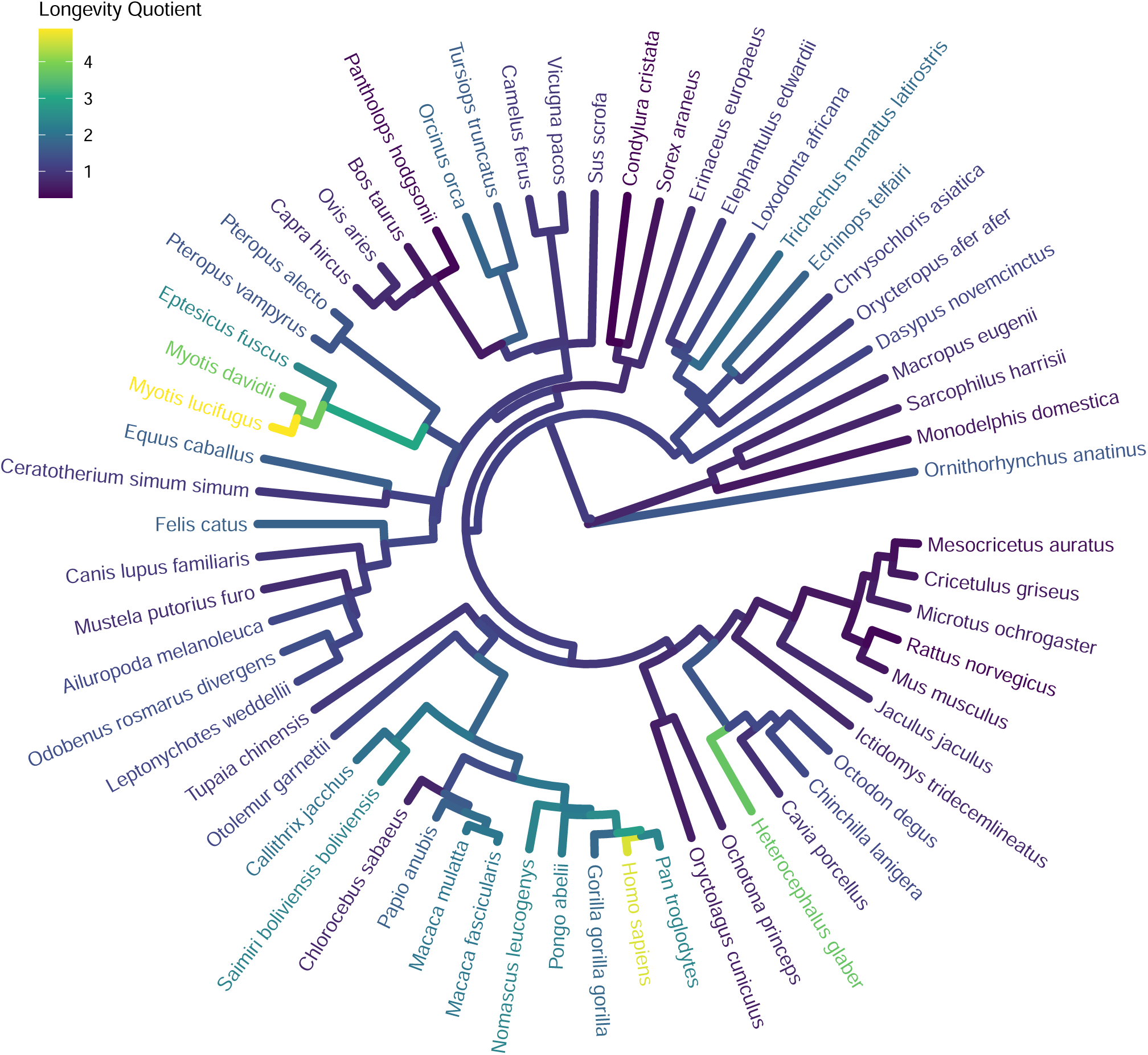
Phylogeny of mammals from [Farré et al., 2021]. Branches are annotated with values of the longevity quotient calculated from Coevol estimates of body mass and longevity (see section 4.8).

In this benchmark we compared our two continuous models to simpler alternatives. First we re-implemented the criterion used in [Farré et al., 2021] to screen sites for potential candidates, namely testing if the 6 longest-lived species had the same exact amino acid and this amino acid was different from the amino acids observed for the 6 shortest-lived species. This criterion is called CAAS (for Convergent Amino Acid Substitution) Scenario 2 in the original publication. A more stringent criterion is also used, where it is asked that the 6 shortest-lived species also have the same amino acid (CAAS Scenario 1). Second, we pre-processed trait values and discretized them into two or three categories, and used the discrete mode of Pelican. We name these approaches `D2 median` and `D3 terciles` respectively. Finally we added a simple ANOVA test, where we use a simple (non-phylogenetic) Gaussian model to predict the longevity quotient of a species from the amino acid it has at the considered site.

Figure 4 shows for each method how recall changes for a given false positive rate (FPR). Model *C_S_* and *C_L_* perform similarly and have the best global performance, particularly for low FPR. Overall, their *p*-values are strongly correlated under *H_A_*(sup. fig. 8). The discretized versions perform worse than both continuous models. This is expected because the discretization is based on trait values at the leaves, but in our simulations, the inflexion point of the sigmoid function for a site is chosen uniformly between the minimum and maximum trait value (see Methods). Thus, it can be far from discretization points, which are chosen using the quantiles of the trait values. As a consequence the discretization is often inadequate to capture the relation between amino acid preferences and trait values. The simple criteria used to define CAAS Scenario 1 and 2 in [Farré et al., 2021] display performances similar to Pelican discretized’s versions on our simulation. More surprisingly, the simple ANOVA test performs similarly to continuous phylogenetic models for FPR greater than 0.01 but its performance degrades for more stringent settings. As a safety check, we performed similar simulations on trees with a different topology, with strong phylogenetic structure, specially crafted to confuse non-phylogenetic methods. More specifically we report in supplementary figure 5 the recall of ANOVA and continuous phylogenetic models at 1% FPR for such trees with an increasing number of leaves. We observe that phylogenetic models display, as expected, increasing recall while it is decreasing for ANOVA. ANOVA is therefore increasingly misled by phylogenetic structure as the amount of data increases, whereas the phylogenetic approaches benefit from the increased amount of information.

**Figure 4:**
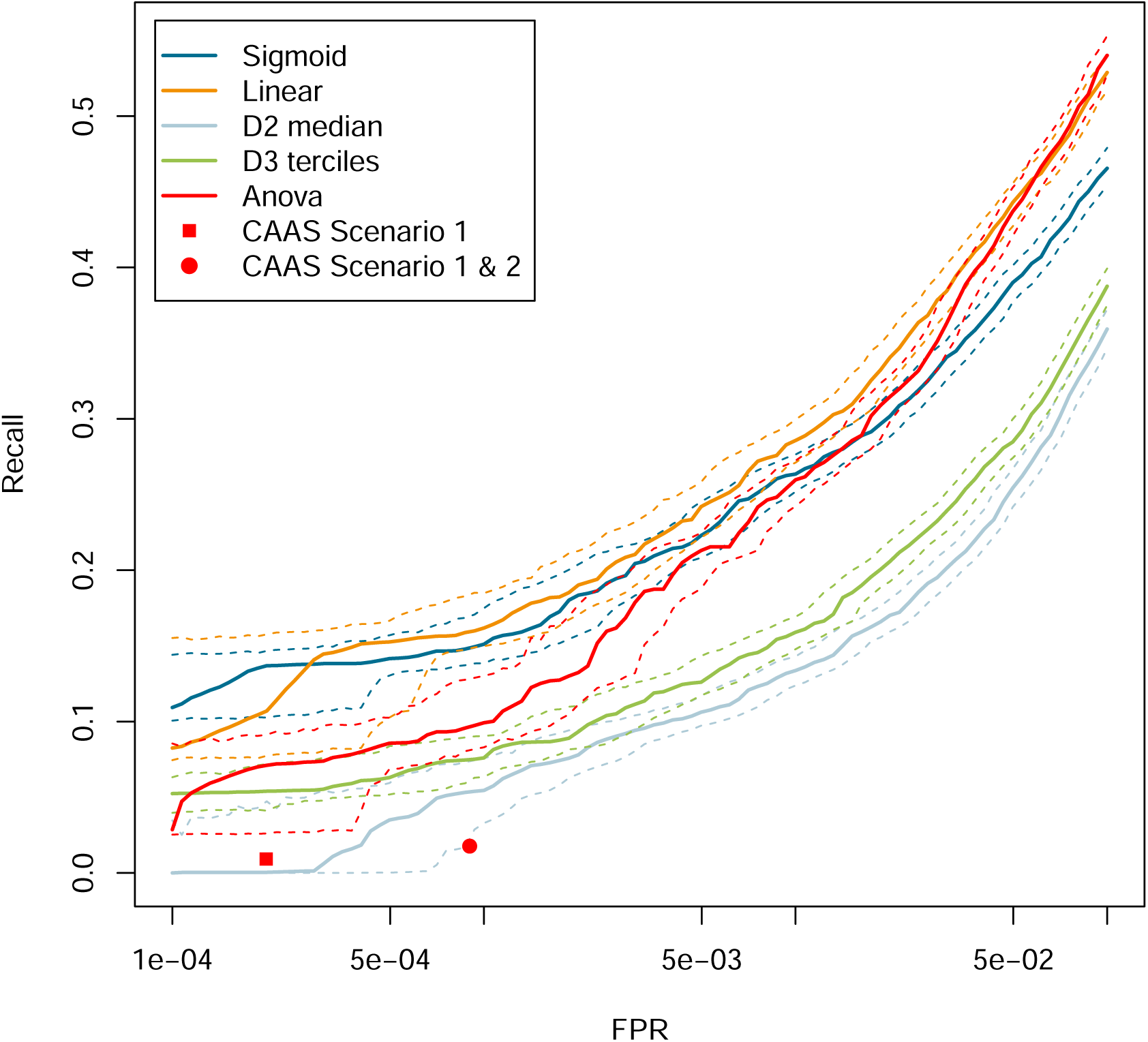
Continuous models *C_S_* and *C_L_* perform better than existing approaches on simulated alignments. On the y axis, recall is displayed as the False Positive rate (FPR) is varied from 0.01% to 10%. Dashed lines show bootstrap 95% intervals, obtained from 1000 replicate data sets, obtained by resampling with replacement the original 20, 000 sites. Red points indicate the performance obtained by CAAS Scenarios from [Farré et al., 2021] method.

The magnitude of the difference in recall values on the mammal phylogeny between continuous phylogenetic models and the other methods suggests a significant improvement useful for genomic scans (fig. 4). However, it is uncertain whether our simulations of *H_A_* sites are realistic because effect sizes for fitness variations have yet to be characterized. Furthermore we use a sigmoid interpolation between asymptotic amino acid frequencies, which might not accurately depict their relation with trait values. Sites simulated under *H*_0_ should be more realistic as we tuned simulation parameters to mirror the amino acid diversity per site in empirical data. Also it is striking that a stringent detection criterion like CAAS Scenario 1 displays a false positive rate on the order of 10*^−^*^4^, which would incur a high number of false positives when run on a genome-wide screen, where millions of sites need to be screened.

### Model *C_L_* suggests limited evidence for previously proposed candidates

We sought to investigate the behavior of model *C_L_* on empirical data, focusing on a few candidates highlighted in [Farré et al., 2021] (corresponding results are shown for model *C_S_* in supplementary figure 6 and 9). [Farré et al., 2021] analyzed thousands of gene families to look for protein coding sites associated with variation in normalized lifespan across mammalian species. With this approach, they notably identified that three genes, WRN, ZC3HC1, and CASP10, previously linked with lifespan, contained 2, 1 and 6 sites of interest, respectively. We reanalyzed these three gene alignments, using the same lifespan data. We used the continuous *C_L_* model on two types of ancestral reconstructions: either using Brownian motion as implemented in Pelican, or based on Coevol [Lartillot and Poujol, 2011]. We ran Coevol on three different concatenates, yielding three separate ancestral reconstructions. Supplementary Figure 7 shows that the three reconstructions provide similar values for the ancestral nodes.

Fig. 5 shows the results obtained with model *C_L_* for all sites of WRN, ZC3HC1 and CASP10, as well as AMY1A that we chose to serve as a negative control. Model *C_L_* was run with the 4 reconstructions of the longevity quotient and we highlight sites such that the mean of the 4 *p*-values is lower than the inverse of the length of the alignment. This threshold is such that we expect that one site per alignment should be highlighted on average, if none is truly associated to the longevity phenotype. However, since the LRT produces *p*-values biased towards low values on this phylogeny, we should actually observe more than one highlighted site on average even if no site is truly associated with longevity.

**Figure 5:**
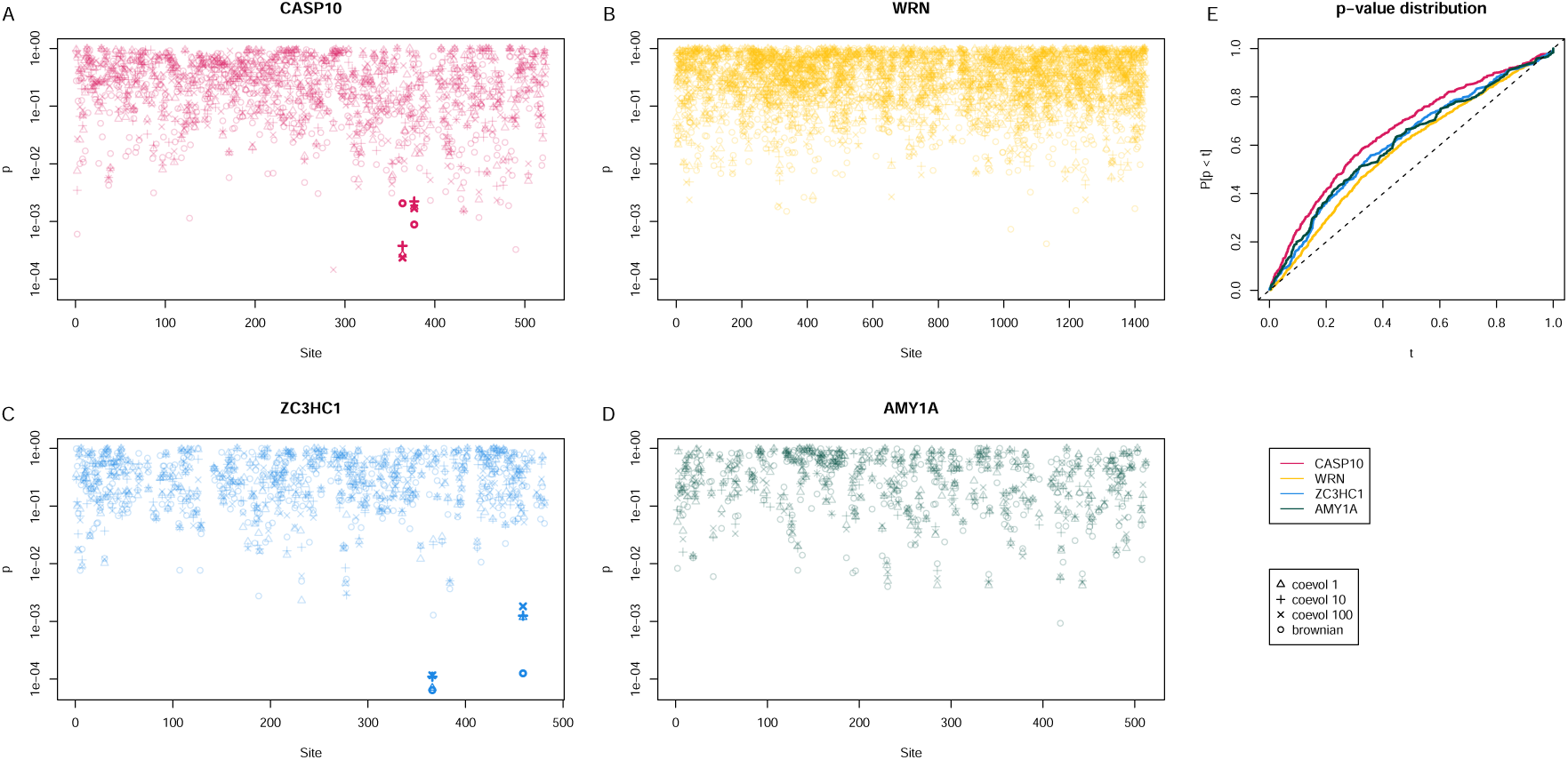
*p*-values obtained with model *C_L_* at each site of genes CASP10 (A), WRN (B), ZC3HC1 (C) and AMY1A (D). At each site, 4 *p*-values are represented, corresponding to 4 different ancestral trait reconstructions. For each site, we compute the mean of the 4 *p*-values and compare this value with the inverse of the alignment length. Sites above this threshold have a more transparent color. E: Distribution of *p*-values for each gene, according to the Coevol 1 ancestral reconstruction.

The protein CASP10 has two highlighted sites (fig. 5, panel A). As noted in [Farré et al., 2021], they are located in the p12 subunit of CASP10 [Cohen, 1997], which starts at position 372 of the protein (which corresponds to approximately the same position in the alignment). This domain is cleaved as part of the activation of the enzyme. It features a higher concentration of lower *p*-values. ZC3HC1 has 2 candidate sites while WRN has none (like AMY1A). We report in Table 1 our results on all sites detected by [Farré et al., 2021], among which the site at position 366 of ZC3HC1 which is the most significant in all three proteins.

**Table 1:** p-values of sites of interest in all three genes WRN, ZC3HC1, and CASP10.

| Gene | Position | Coevol1 | Coevol10 | Coevol100 | Brownian |
|---|---|---|---|---|---|
| WRN | 1018 | 0.018 | 0.011 | 0.011 | 0.014 |
| WRN | 1055 | 0.077 | 0.17 | 0.1 | 0.014 |
| ZC3HC1 | 366 | 7 × 10<sup>−5</sup> | 0.00011 | 0.00012 | 6.4 × 10<sup>−5</sup> |
| CASP10 | 363 | 0.12 | 0.074 | 0.1 | 0.082 |
| CASP10 | 370 | 0.16 | 0.15 | 0.14 | 0.023 |
| CASP10 | 371 | 0.14 | 0.19 | 0.23 | 0.07 |
| CASP10 | 412 | 0.019 | 0.021 | 0.029 | 0.0033 |
| CASP10 | 440 | 0.061 | 0.084 | 0.099 | 0.023 |
| CASP10 | 464 | 0.04 | 0.055 | 0.036 | 0.032 |

The *p*-values we find on the 3 alignments are only mildly significant at best: only the 2 highlighted sites in ZC3HC1 and 10 in CASP10 are significant at 5% FDR according to a Benjamini-Hochberg estimate; but again since *p*-values are biased towards low values, those FDR estimates probably are optimistic. Indeed the distribution of empirical *p*-values shown in fig. 5 Panel E is similar to what we obtained on simulations, although we observe CASP10 has a markedly different distribution from the 3 other proteins. According to our model there is thus limited evidence in the sequence of the 3 genes for an association with the longevity quotient. That does not preclude a true relationship exists and independent assays can be used to validate the most promising sites, as was done in [Farré et al., 2021].

### Detailed phylogenetic analysis of candidate sites

The site at position 1018 of WRN complies with CAAS Scenario 1: all 6 shortest-lived species have a L at this position while all longest-lived species have a F. It was at first puzzling to us why it is not considered particularly significant by our continuous models. A first answer is illustrated by Fig. 4: it is not very rare to observe sites complying with CAAS Scenario 1 under the null hypothesis of no association with the phenotype. We sought to gain more insight in the matter by observing the detailed history of sites with respect to the evolution of the phenotype. To this end, we computed substitution mappings using the algorithms developed in [Rodrigue et al., 2008], and we generated 100 detailed substitution histories for the site assuming the homogeneous model. In parallel we fit model *C_S_* on the site to estimate the two asymptotic profiles. We then report the distribution of trait values where the substitution occurs, distinguishing between transitions that are favored in the low-end profile of the sigmoid function and those that are favored in the high-end profile. In figure 6, we show the result for three sites: WRN:1018, CASP10:464 and ZC3HC1:366. For WRN:1018 (top left part of the figure), the two histograms are nearly completely overlapping, which means that transitions towards F in general did not happen on branches with higher longevity values than transitions towards L. This result is consistent with previous findings by Farré et al., where WRN:1018 was identified but not validated (*p* = 0.63). On the contrary, we see more contrast between the blue and orange histograms for CASP10:464 and ZC3HC1:366, whose *p*-values are smaller by 2 to 4 orders of magnitude. In [Farré et al., 2021], these two substitutions were consistently validated, with *p*-values of 0.00278 and 0.00418, respectively.

**Figure 6:**
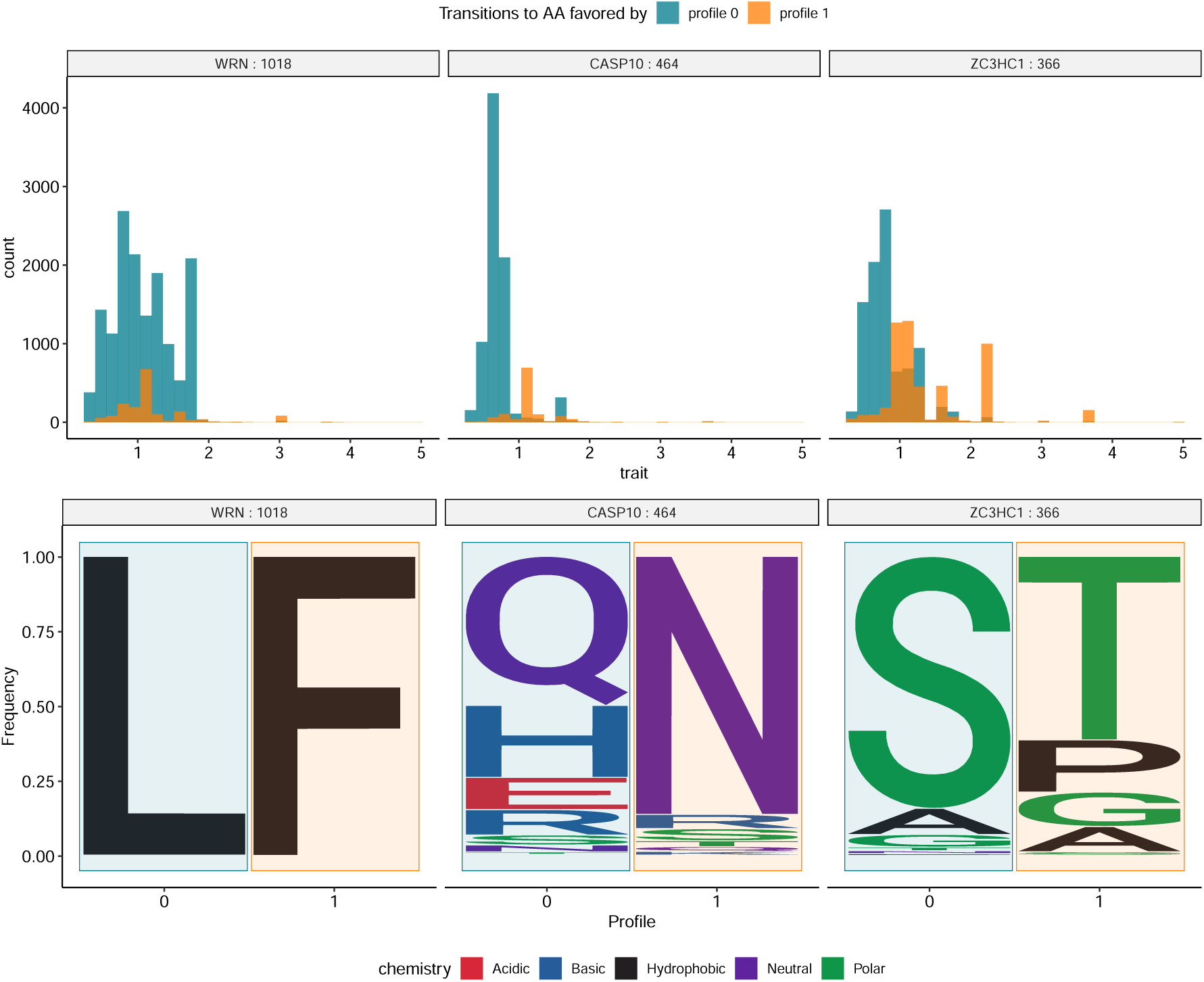
Detailed analysis of longevity-amino acid substitutions relationship in 3 sites. (Lower part) Asymptotic profiles estimated using model *C_S_* on each site. (Upper part) Histograms of trait values at each substitution in 100 substitution mapping histories under the homogeneous model. Substitutions are separated in two groups depending on the target amino acid: values associated to transitions towards an amino acid favored in low-end (resp. high-end) profile are depicted in blue (resp. in orange).

## 3 Discussion

Our models *C_S_* and *C_L_* to study associations between protein sites and continuous phenotypes across species condition protein site evolution on a particular phenotypic evolutionary history, where each branch of the phylogeny is annotated with a phenotypic value. It is convenient to assume that phenotype evolution is known, but in practice it may not be easy to reconstruct ancestral phenotypic values, and therefore errors could be made. A standard approach to circumvent this difficulty would be to jointly reconstruct protein site and phenotype along the phylogeny, allowing to integrate over the uncertainty in phenotype evolution when studying protein site evolution. However, in our case, this approach is problematic because we reason one individual site at a time, when phenotype evolution should be shared across all sites. Therefore we chose to study protein site evolution conditionally on phenotypic evolution, in keeping with model *D*. Models *C_S_* and *C_L_* have been implemented within Pelican, and build upon the same underlying model of sequence evolution as model *D*. Model *D* was proposed in [Tamuri et al., 2009b], and has been shown to have the best power in a benchmark of 6 methods to detect sites associated to a discrete phenotype, with a computational footprint lower than other methods that rely on phylogenetic models of sequence evolution [Duchemin et al., 2022]. In particular, although it relies on protein sequences, its performance was similar to or better than methods that analyze codon sequences. We therefore expect that little gain would be obtained by using codon sequences instead of protein sequences for models *C_S_* or *C_L_*.

We evaluated the performance of the two new models to detect sites associated with a continuous phenotype on data simulated along a phylogeny of 62 mammals, annotated with the longevity quotient of the species included in the data set [Farré et al., 2021]. This provided us with a realistic continuous variable at the tips of the phylogeny. Our simulations show that model *C_L_* and model *C_S_* have better performance for low false positive rates than simpler alternatives, model *C_L_* performing slightly better than *C_S_*. We believe this is because *C_L_* achieves a better trade-off between number of parameters and fit to the data than the *C_S_* model. The two additional slope and shift parameters of the *C_S_* model do not enable it to significantly better describe *H_A_* site evolution but raise the expectation in likelihood difference between the heterogeneous and the homogeneous models. We also noted that a simple ANOVA performs well on the phylogeny we tested, and at much lower computational footprint than model *C_L_* and *C_S_*. However we showed that it may also perform significantly worse on a topology where some leaves are very correlated with each other. In such a case, considering that all leaves are independent leads to wrong inferences, and increasing the size of the topology actually worsens the prediction performance. We suggest that it is still useful to use an ANOVA test for filtering out unpromising candidates; in practice setting a p-value threshold on the order of 10*^−^*^3^ is enough to efficiently and safely discard a large portion of the input sites. Both continuous models outperform discretized models working with categories of longevity quotients. They also outperform the use of CAAS proposed in [Farré et al., 2021]. As a result, we expect that the new models, model *C_L_* in particular, could become very useful to researchers studying associations between continuous variables and protein sequences. We now comment on a few limitations and perspectives of this work.

The results of our simulated benchmark, as well as the detailed analysis of substitution histories we showed on a few sites, confirm the importance of modeling the evolutionary path of genotype and phenotype instead of simply focusing on the amino acid states at the tips. However, this assumes that it is possible to reconstruct past phenotypic states with confidence. In our simulation setting we assumed a perfect knowledge of the phenotype in all species of the tree. Ancestral phenotype reconstruction is however a challenging task and the statistical power of our continuous models would be to some extent lower had we introduced some noise in the trait values. On empirical data we observe that calculated *p*-values sometimes vary by one or two orders of magnitude between equivalent but distinct reconstructions of the phenotype (see figure 5 and table 1), although most of the time the impact is limited. A useful enhancement of our method would be to incorporate some representation of the uncertainty on ancestral phenotypic values.

Our simulations were sized according to the dataset of [Farré et al., 2021], which contains 62 mammal species. Our results show that even the best method we considered has a very low recall when set to a false positive rate compatible with genomic scans. We surmise that this is primarily due to the sample size, which is too low to properly estimate the parameters of the site-specific models. A lot more mammal proteomes are now available and a similar analysis could be performed with more than 400 species. It is also possible that increasing the sample size could improve calibration substantially.

Model *D*, *C_L_* and *C_S_* are designed to estimate how amino acid preferences vary with a covariable. That makes them better suited to detect changes in the direction of selection than changes in its intensity, even though they can account for varying rates of evolution. Variations of amino acid preferences may have two different causes: first, they could stem from a change in fitness, *i.e.* a change in the direction of selective pressure; second, it could be that the fitnesses are unchanged but the intensity of selection has varied. The second case happens when the effective population size of a species varies, while the first case corresponds to a *bona fide* adaptation to a changing environment. An important question is thus to distinguish between these two situations, and that is imperfectly handled as of today. This is made more difficult in the particular case of our empirical dataset, since longevity is known to be negatively correlated with effective population size. We suspect that this adds some noise in this analysis, as the reduced population size affects the intensity of selection on all sites, among which some of them would also undergo changes in the direction of selection. For instance, the accumulation of mildly significant sites in the p12 subunit of CASP10 could be interpreted as an increased selective pressure on this domain in long-lived species, rather than a specific adaptation for this phenotype. Indeed all sites reported by [Farré et al., 2021] in CASP10 correspond to CAAS Scenario 2. With model *D*, *C_L_* and *C_S_*, it is possible to inspect the estimated profiles and get a sense of the evolutionary scenario at hand. If we have a look at the estimated profiles for site 366 in ZC3HC1 (figure 6), we notice that they include both hydrophobic and polar amino acids, and that the main amino acids in each profile, Serine and Threonine, are fairly similar. Based on these considerations, this site does not seem to have undergone a very significant change in the direction of selection. However, a testing procedure to distinguish changes in the direction of selective pressure from changes in selection intensity would help significantly for the study of parallel adaptations.

An important limitation of our approach is that the prerequisites to apply the likelihood ratio test we use are not met, resulting in a non uniform distribution under *H*_0_ and imperfect calibration of the test. This implies that a reliable significance threshold might be very dependent on each specific dataset, especially when working with a lower number of sequences in the alignment. In practice, this can be circumvented by either: (1) using simulations under *H*_0_ to calibrate the test (computationally intensive); (2) ranking *p*-values to identify the best candidate sites for genotype-phenotype association in the alignment.

Overall, we have introduced new models to search for protein sites associated with continuous traits along a phylogeny. After evaluating their power, we concluded that they will be most useful on datasets containing large numbers of sequences, which are becoming more common. Further development will focus on refining the detection of various selective regimes, notably to contrast changes in the intensity of selection from changes in the direction of selection.

## 4 Methods

### 4.1 Models of sequence evolution

We use a model of codon sequence evolution to simulate sequences, and a model of protein sequence evolution for inference and for checking the implementation. In both cases, the evolution of each site is modeled independently of the other sites. Both models are continuous time Markov processes running along the branches of a phylogeny whose branch lengths are in units of expected number of substitutions [Felsenstein, 1981]. At each branch *b*, substitution rates are described by an instantaneous substitution matrix *Q_b_* which, combined with a branch length *l_b_*, is used to compute a substitution probability matrix *P_b_* as follows:

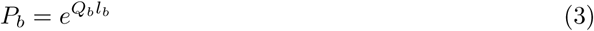

Individual sites are associated to specific amino acid preference parameters to account for heterogeneity in the process of evolution across sites. Different amino acid preferences may also be used across branches, if the site is assumed to be associated to the continuous phenotype under consideration. Such sites are called *H_A_* sites, whereas *H*_0_ sites are not associated to the continuous phenotype. As a result, for *H*_0_ sites, *Q_b_* = *Q* for all branches *b*.

### 4.2 Model of codon sequence evolution for simulations

We use a mutation-selection model of codon sequence evolution [Halpern and Bruno, 1998, Yang and Nielsen, 2008, Rodrigue et al., 2010, Tamuri et al., 2012, Bloom, 2014] to simulate sequence evolution using the Gillespie algorithm [Gillespie, 1976]. Mutation-selection models handle mutations at the DNA level, and selection at the amino acid level. Both are combined into a 61 × 61 codon substitution matrix. A matrix of mutation probabilities *µ* between individual nucleotides is derived from exchangeabilities and equilibrium frequencies [Lanave et al., 1984, Tavaré et al., 1986]. Exchangeabilities are drawn from Gamma(1, 1) distributions, and equilibrium frequencies from a Dirichlet(10, 10, 10, 10) distribution, and are shared across sites. A 61 × 61 codon mutation matrix is built from this nucleotide mutation matrix by considering that double and triple substitutions between codons are not allowed.

Selection coefficients 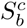 (eq. 4) associated to the amino acids are defined as the difference in scaled fitness *F* between the ancestral amino acid *X* and the descendant amino acid *Y*, according to the parameters applying on the current branch *b*. The relative fixation probability 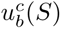 for a mutation at site *c* and branch *b* is computed from the selection coefficient *S* as per [Kimura, 1983]:

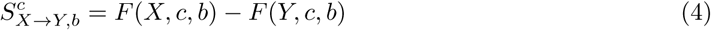

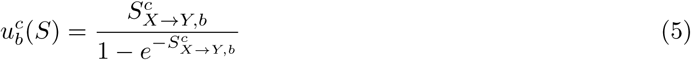

Fitness values are drawn from a set of 263 preset frequency profiles [Rey et al., 2019] for each site. These frequency profiles are transformed into fitness profiles by calculating their logarithm and multiplying the result by a factor *Ne* = 2. One profile needs to be drawn for *H*_0_ sites, whereas more parameters need to be drawn when simulating *H_A_* sites (see section 4.4). Codon substitution rates *Q_b_* are the product of mutation rates *µ* and the relative probability of fixation :

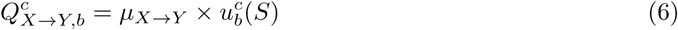

### 4.3 Model of protein sequence evolution for inference

For *H*_0_ sites, the model used for inference in Pelican is similar to commonly used models of protein sequence evolution [Whelan and Goldman, 2001, Le and Gascuel, 2008]. In particular, it can be expressed as a combination of a 20 × 20 exchangeability matrix and a vector of 20 equilibrium amino acid frequencies. For exchangeabilities we use the values from the WAG matrix [Whelan and Goldman, 2001]. Site-specific equilibrium frequencies are estimated in the maximum likelihood framework for the *H*_0_ and D models, or are computed with a sigmoid or log linear function that links phenotypic value and amino acid preference as explained below.

### 4.4 Functions used to link phenotypic value and amino acid preference

In *H_A_* sites, we model amino acid preference at a site as depending on a continuous phenotypic trait. In a mutation-selection framework operating at the codon level, this results in different amino acid fitnesses depending on the phenotypic trait value. In a model of protein sequence evolution, this results in different amino acid frequencies depending on the phenotypic trait value. We use the mutation-selection framework for simulating sequences, and rely on the model of protein sequence evolution for inference inside Pelican.

For all variants of our model, the instantaneous transition rate *Q_ij_* between two distinct states *i* and *j* on a particular branch can be expressed as

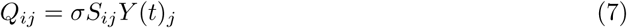

here *σ* is a scaling factor accounting for variations in the overall rate of evolution, *S* is the WAG matrix and *Y* (*t*) is the vector function used to link phenotypic value *t* to amino-acid preferences. Diagonal elements of the rate matrix *Q* are by definition:

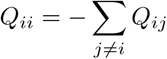

At a given branch, the value of *t* is the trait value provided as input to the method for the descendant node of the branch. These values are user-input, and typically correspond to trait measurements at the tips, and to ancestral trait reconstructions for internal nodes. We provide basic ancestral trait reconstruction through Brownian motion if users cannot rely on a better approach. We now compare two different specifications of *Y* (*t*) : one is a sigmoid (defining the model we call *C_S_*, the other is a multinomial logistic regression (defining the model we call *C_L_*).

#### 4.4.1 Sigmoid function

As a first alternative, we model amino acid preferences (expressed as fitness values or frequencies) at a particular site as a smooth interpolation between two asymptotes. For this we use a sigmoid vector function *Y* of the trait value *t*:

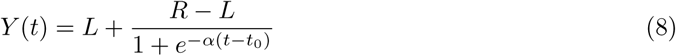

where *L* and *R* are left and right asymptote respectively, *t*_0_ is the trait value at the inflection point of the sigmoid and *α* a slope. See figure 2 for an illustration of the general shape of the function. When modeling frequencies, the entries of *L* and *R* are positive and sum to one, and so does *Y* (*t*). The left and right asymptotes require a total of 40 parameters for fitnesses and 38 for frequencies (since frequencies sum to 1 in each vector). In practice, we set the frequencies of unobserved amino acids to 0.0 as in [Tamuri et al., 2009b, Duchemin et al., 2022], which strongly reduces the number of parameters (see section 4.5). The slope and inflection point parameters that control the shape of the sigmoid are shared across amino acids, adding two parameters in the model.

#### 4.4.2 Multinomial logistic regression

The second alternative to model amino acid frequencies as a function of phenotypic value is based on the combination of a linear function and a softmax link function, as in multinomial regression models :

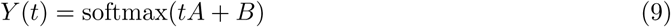

where *A* and *B* are two real vectors of size 20. Since softmax(*X* + *α***1**) = softmax(*X*) for any real value *α*, we can without loss of generality enforce that the input vector to the softmax function sums to 0. One way to achieve that is to use vectors A and B of size 19, compute *tA* + *B* and add a 20th coordinate to it so that the sum of all coefficients equals 0. As in the sigmoid model, frequencies of unobserved amino acids are set to 0. Overall, this formulation uses two fewer parameters compared to the sigmoid model. Note that its asymptote for large (resp. small) trait values is a Dirac distribution on the largest (resp. smallest) entry in *A*. This is illustrated on figure 2: while the multinomial logistic model can be adjusted to be close to the sigmoid model on a given interval (red rectangle in the figure), it doesn’t have the same behavior when evaluated outside of this interval. Most notably, some amino acid may have a non monotonous dependence to the trait value.

#### 4.4.3 Models of sequence and continuous trait evolution

The model of amino acid sequence evolution described above (section 4.3) is combined with the sigmoid (section 4.4.1) or multinomial logistic models (section 4.4.2) to link trait and site evolution to give the *C_S_* and *C_L_* models. These models also include a scale parameter that must be estimated, and that corresponds to a site-specific rate. Fixing the scale parameter to 1.0 resulted in lower detection power (not shown). The *C_S_* and *C_L_* models are implemented in Pelican and can be used for inference.

### 4.5 Sparse parameter estimation at each site

At each site, not all 20 amino acids are observed. Except for particular exchangeability matrices this leads to maximum likelihood estimates of 0 for the equilibrium frequency of these amino acids. In order to reduce the number of parameters to consider during inference, we only work with the subset of amino acids that are observed at the site. Therefore, during estimation, fewer parameters need to be estimated per site. For instance, for a site that has only 5 different amino acids, model *C_S_* includes only 11 parameters and model *C_L_* 9 parameters instead of 41 or 39 if all 20 amino acids are considered. This heuristic improves the speed of numerical calculations, but also provides a better (although not perfect) estimate of the effective number of degrees of freedom in the likelihood ratio test (see below).

### 4.6 Likelihood ratio test in continuous models

The discretized version of Pelican relies on a likelihood ratio test between a homogeneous model, where the same parameters apply throughout the tree, and a model where different amino acid equilibrium frequencies apply depending on the condition associated with the branch [Duchemin et al., 2022]. The distribution of the likelihood ratio under the null hypothesis is unfortunately unknown and we applied the same heuristic than in [Tamuri et al., 2009a]. We adopt the same framework to assign *p*-values to sites, because the homogeneous model is also nested within models *C_S_* and *C_L_*. For *C_S_*, the number of parameters that need to be considered in this test is the two parameters in the sigmoid function, plus the number of amino acids present at the site, minus one. For *C_L_*, the number of parameters that need to be considered in this test is the number of amino acids present at the site minus one. For a site where 4 amino acids are observed, the likelihood ratio test (LRT) is computed as follows. First, compute the log-ratio *LR*:

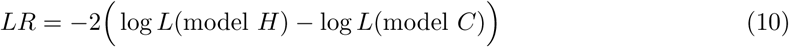

where model *H* is the homogeneous model and model *C* is either *C_S_* or *C_L_*. Then compare *LR* to a *χ*^2^ distribution with 4 *−* 1 + 2 = 5 degrees of freedom for *C_S_* and 3 degrees of freedom for *C_L_*.

### 4.7 Benchmark simulations

#### 4.7.1 Tree and trait values

We used data from [Farré et al., 2021]. The phylogeny of mammals contains 62 species, and we used longevity quotient values to annotate traits at the tips. Ancestral trait values at internal nodes were estimated either using Brownian motion, or using Coevol [Lartillot and Delsuc, 2012] on three gene concatenates (see section 4.8). Figure 3 represents the phylogeny annotated with Coevol trait values obtained with the first concatenate.

#### 4.7.2 Simulations

We performed two types of simulations.

##### Simulations to check the implementation

The first type of simulation was based on the model used for inference. These simulations were used only to check that the inferred parameters converge towards the true values as the amount of data increases. To do so, 70 sites were simulated along balanced, symmetric phylogenies of sizes ranging from 10 to 1,000 leaves. Amino acid profiles were sampled in a Dirichlet distribution with concentration parameters equal to 0.1. Supplementary figure 1 shows that parameters of the sigmoid models were found to converge to their true values as the size increases, suggesting that the optimization routines used in our software are well behaved.

##### Simulations to evaluate the models

The second type of simulations were performed in a manner similar to [Duchemin et al., 2022], but adapted to continuous phenotypes. More precisely, simulations were based on a mutation-selection model, where each site is associated to site-specific amino acid fitness profiles. Two types of sites were simulated: *H*_0_ sites not associated to the phenotypic trait, and *H_A_* sites associated to the phenotypic trait. The former were simulated under a constant mutation-selection model applying to all branches of the phylogeny. A single vector of amino acid fitnesses was drawn as in [Duchemin et al., 2022] and applied throughout the tree. The latter were simulated under mutation-selection models that changed at each branch of the phylogeny. More precisely, equilibrium amino acid frequencies changed depending on the phenotype value at the tip of the branch. To this end, 2 vectors of amino acid fitnesses were drawn per site, and were used to provide left (*L* in Equation 8) and right (*R* in Equation 8) asymptotic amino acid fitness values, for low and high values of the continuous phenotype respectively. Then, at each branch, the end phenotypic value was used to compute branch-specific amino acid fitnesses, according to the sigmoid link function between phenotype and amino acid frequency (figure 2, Equation 8). The shift parameter *t*_0_ and the slope parameter *α* of the sigmoid are simulated using the *t*_min_ and *t*_max_ extremal values of the trait :

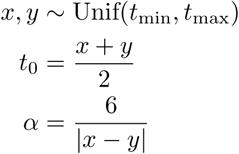

We checked that the simulated sites were similar to empirical sites, by computing site-wise amino acid diversity (the number of different amino acids observed at a site) in both types of data sets. To this end, we used 3 alignments from [Farré et al., 2021], and compared them to 1000 *H*_0_ sites and 1000 *H_A_* sites. Supplementary Figure 3 shows that *H*_0_ sites are similar to sites found in mammalian alignments from [Farré et al., 2021], but *H_A_* sites have a higher diversity. This is expected as *H_A_* sites have been simulated with two amino acid profiles, leading to observe up to twice as many amino acids as in *H_A_* sites simulated with a single profile.

#### 4.7.3 Evaluated methods

We compared the continuous version of Pelican to the approach used in [Farré et al., 2021], and to the discrete version of Pelican, applied to the phenotype after discretization of the phenotypic trait into categories.

[Farré et al., 2021] used a two-step approach to identify candidate sites. First, they looked for sites where all 6 longest-lived species had the exact same amino acid, without gaps. Then among those they selected sites where the amino acids for the 6 shortest-lived species were different from that found in the longest-lived species. Cases where the shortest-lived species share the same amino acid were called CAAS Scenario 1 (where CAAS stands for Convergent Amino Acid Substitution); other cases were called CAAS Scenario 2.

We tried different ways to discretize the trait values in Pelican’s discrete model. Model `D2 median` works with 2 categories of the trait value, based on whether they fall below or above of the median of the trait value observed at the tips. We also considered variants of this where the median is replaced by the mean or the phylogenetic mean (that is, the inferred trait value at the root of the tree, assuming a Brownian model for the trait) with qualitatively similar results (data not shown). Model `D3 terciles` discretizes trait values into 3 categories based on terciles of the trait value observed at the tips. All methods were run on the simulations described above, and their performance was measured using the recall at different rates of false positives. We chose to focus on this metric as it seems to be the most useful when considering genomic screens for sites associated to a phenotype of interest. Indeed, we expect that only a small proportion of sites in the proteome will be associated to the phenotype, which means that even a low false positive rate can overwhelm genuine signal. As a numeric illustration, if we assume that there are about 5 million sites in a proteome, and only 1, 000 sites associated to the phenotype, a false positive rate of 0.1% contributes 5, 000 false positives. This means that the experimental validation of candidates might be successful only for 1 site out of 6.

### 4.8 Inference on empirical data

We used four alignments from [Farré et al., 2021], namely `NM 000553`, `NM 001282190`, `NM 032977` and `NM 004038`, corresponding to the genes WRN, ZC3HC1, CASP10 and AMY1A respectively. We collected the longevity data normalized by weight from the same source, and used model *C_L_* in Pelican to search for sites associated to longevity in the three genes. The model was run on the log-transformed normalized longevity data in two different ways. First, ancestral trait values were inferred with Brownian motion as implemented in Pelican. Second, we ran three inferences based on different ancestral reconstructions of the log-transformed normalized longevity data. These three reconstructions were obtained by running Coevol [Lartillot and Delsuc, 2012] on three different concatenates of nucleotide sequences. For this we proceeded as follows.

We obtained the 13,035 nucleotide alignments screened for association with variation in longevity in mammals from [Farré et al., 2021]. We concatenated them in random groups of 20 alignments, and arbitrarily chose concatenates 1, 10, and 100 for the subsequent steps. We ran two chains of Coevol on 8 processors each, on each of the three concatenates, and using the UCSC database hg38 tree pruned to contain only the 62 species studied in [Farré et al., 2021]. We used the “dS and omega” model (`dsom`) to estimate correlations between log-transformed longevity data, log-transformed weight data, and both dS and dN/dS. We set the root node age to 208.15 Million years ago (Mya) with a standard deviation of 44, and used 19 calibrations obtained from [Lartillot and Delsuc, 2012] and from [Álvarez Carretero et al., 2022]. The calibrations used are show in Supplementary table 1. Chains were run for more than 7,400 iterations. Expected Sample Size values as computed by `tracecomp` were all superior to 100, except for the parameter `root_1` for alignment 10, which was 54. We discarded 2 000 iterations as burnin and estimated median trait values at ancestral nodes of the tree using `readcoevol`. Then we used Python scripts to compute normalized longevity ancestral trait values at each node of the phylogeny, and to write the data as NHX files for input into Pelican.

## Availability

Code for this work is available at https://gitlab.in2p3.fr/phoogle/pelicon-paper and data at https://doi.org/10.5281/zenodo.21788251

## Acknowledgements

We thank Arcadi Navarro Cuartiellas for fruitful discussions.

