## Supplementary information for "Phylogenetic detection of protein sites associated with continuous traits"

March 2026

The Orthomam phylogeny and the body-mass trait annotation for each of its species that we used in our simulations are available as supplementary data at https://doi.org/10.5281/zenodo.7414499.

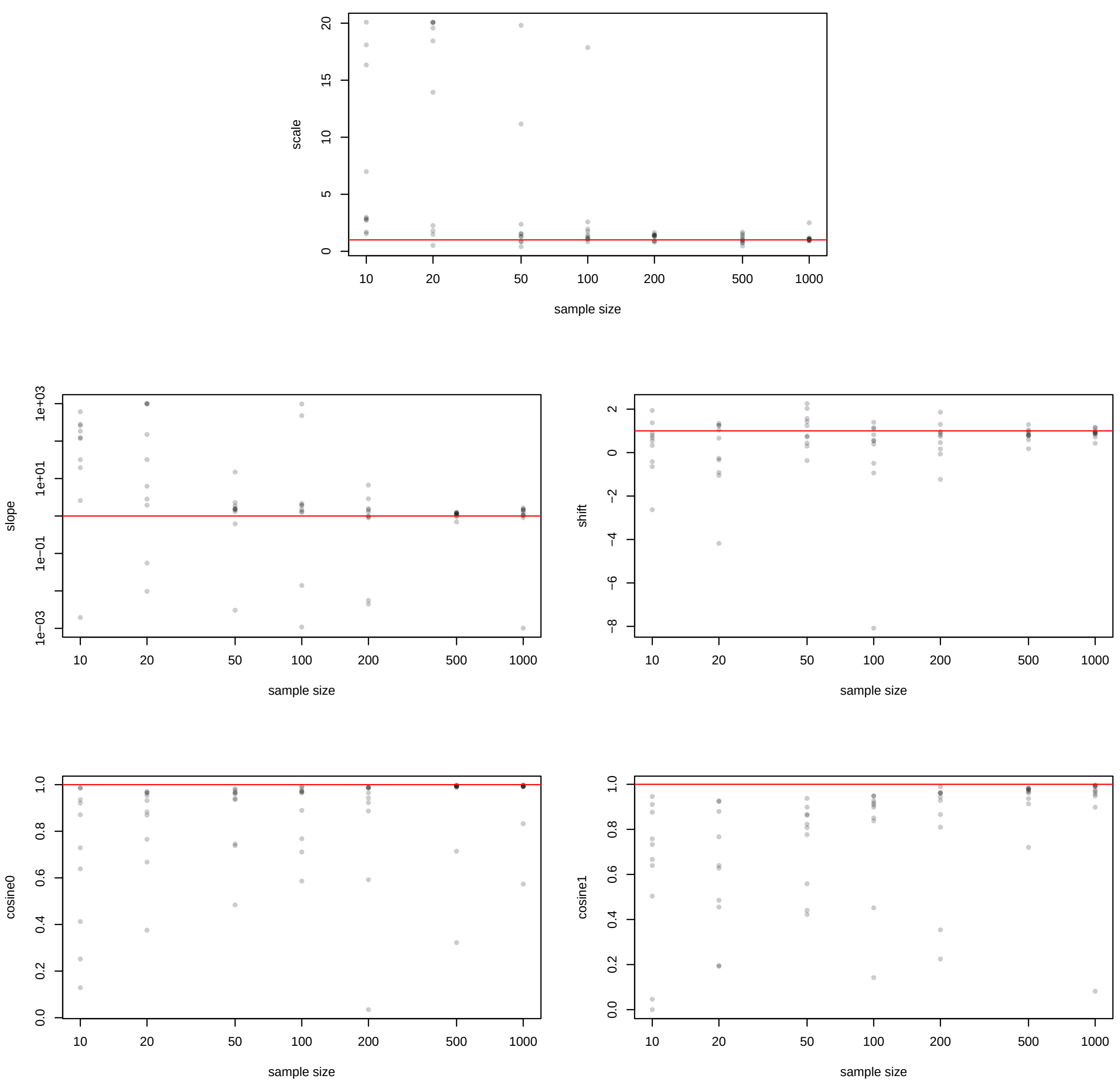


Supplementary figure 1: Estimated parameters of the sigmoid model from simulations on trees of increasing size. Each plot corresponds to a parameter (scale, slope, shift, left and right asymptotes) and displays the estimated value (or its cosine with the true value in the case of vector asymptotes) in Y-axis as a function of sample size in X-axis. For each sample size, 10 points are displayed, each one representing a simulated site. The horizontal red line represents the true value of each parameter.

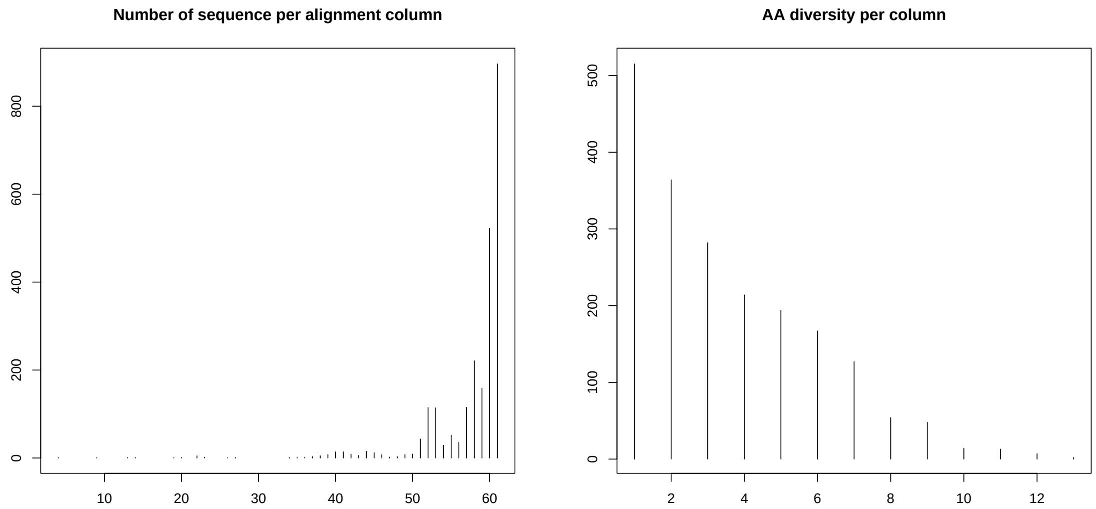


Supplementary figure 2: Summary statistics of WRN, ZC3HC1 and CASP10 alignments. Left: number of species across positions of the alignments. Right: Amino acid diversity across positions of the alignments. Most sites in the alignments have fewer than 6 different amino acids.

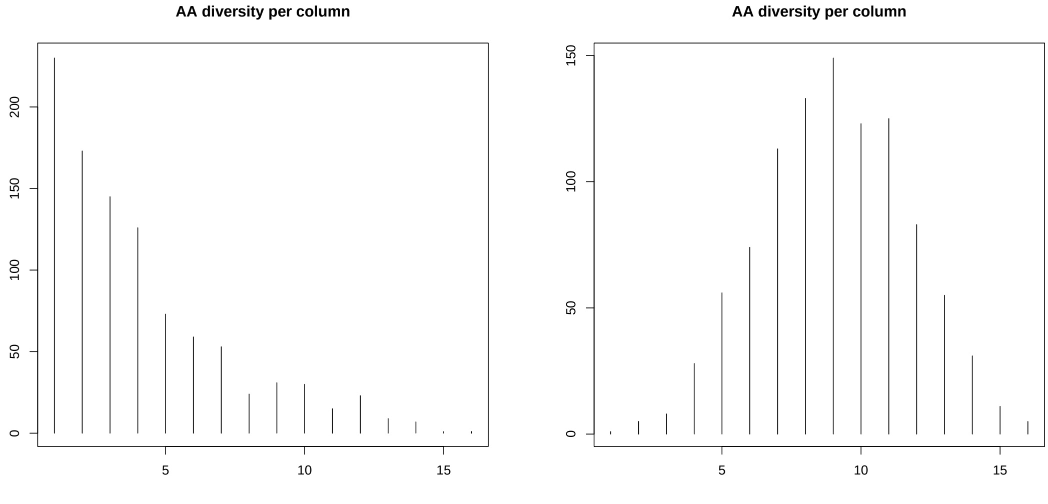


Supplementary figure 3: Amino acid diversity per position in sites simulated with our mutation-selection model. Left: Sites simulated under the homogeneous $H_0$ model. The diversity per site follows a trend similar to that observed in figure 2. Right: Sites simulated under the heterogeneous $H_A$ model. The diversity per site is higher than that observed in most empirical sites.

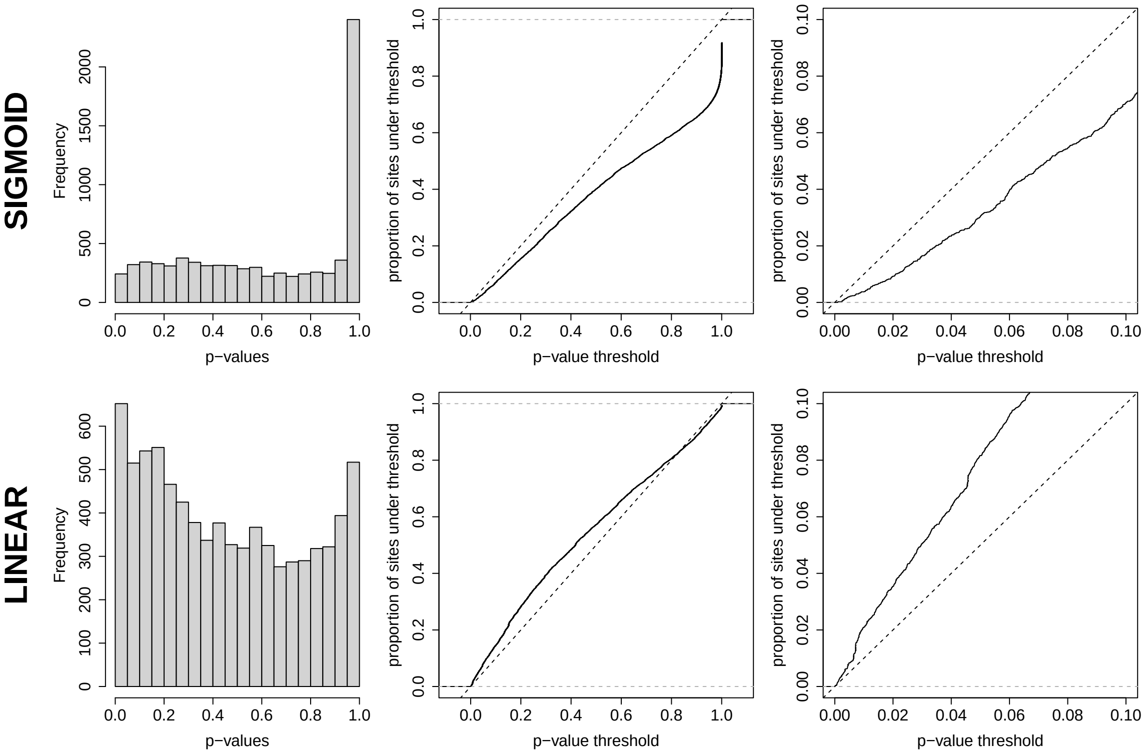


Supplementary figure 4: Distribution of $p$-values calculated using the LRT on model $C_S$ (above) and $C_L$ (below), on $H_0$ sites. Only non constant sites (that is with more than one amino acid at the leaves of the phylogenetic tree) are used. (Left) Histogram representation (Middle) Cumulative distribution function representation (Right) Zoom on the cumulative distribution where $p \leq 0.1$.

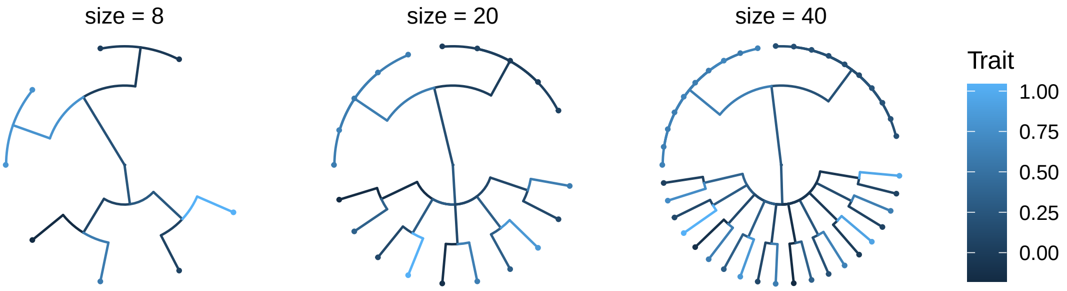


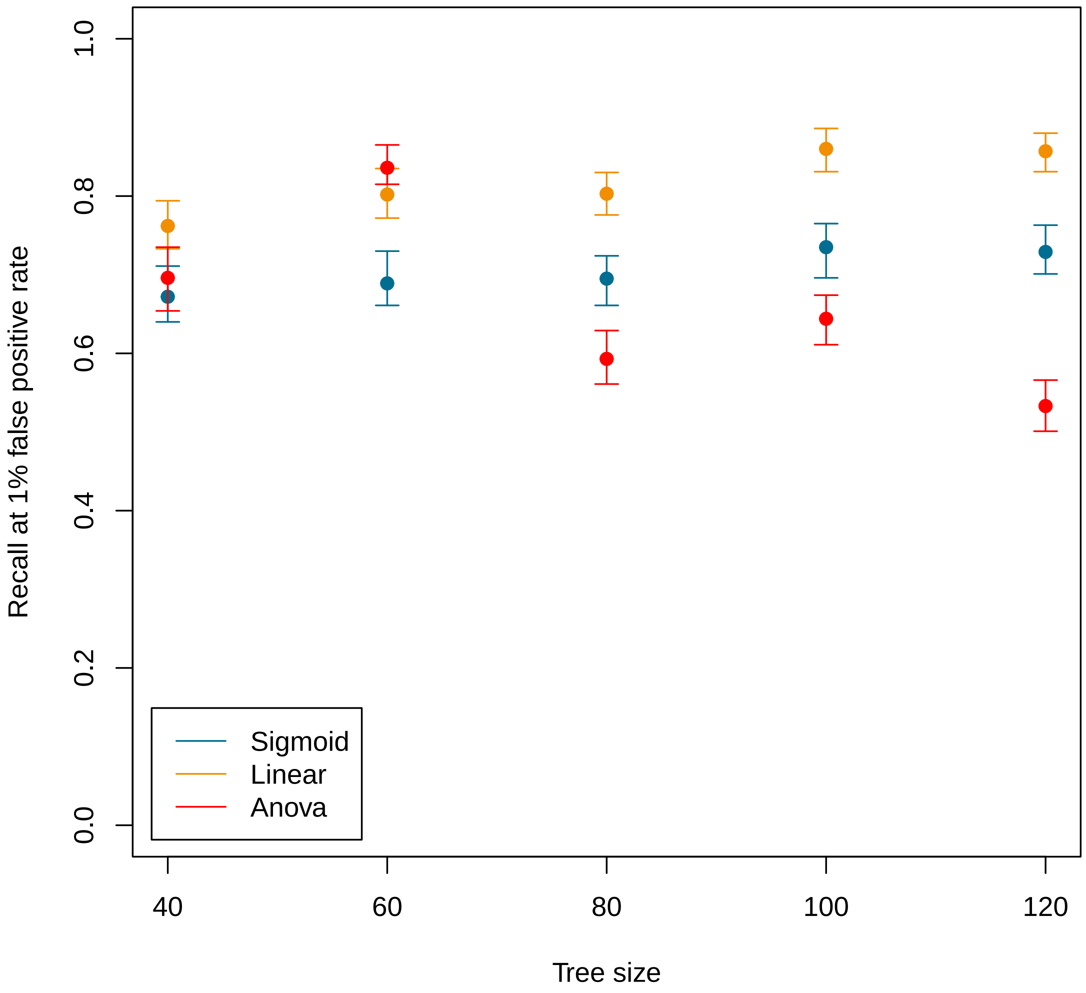


Supplementary figure 5: Second benchmark performed on a topology specially crafted to confuse non-phylogenetic methods. (Top) Synthetic topologies used in the benchmark. (Bottom) Recall at 1% FDR as a function of tree size for model $C_S$, model $C_L$ and a simple ANOVA test. Estimates and 95% confidence intervals are shown.

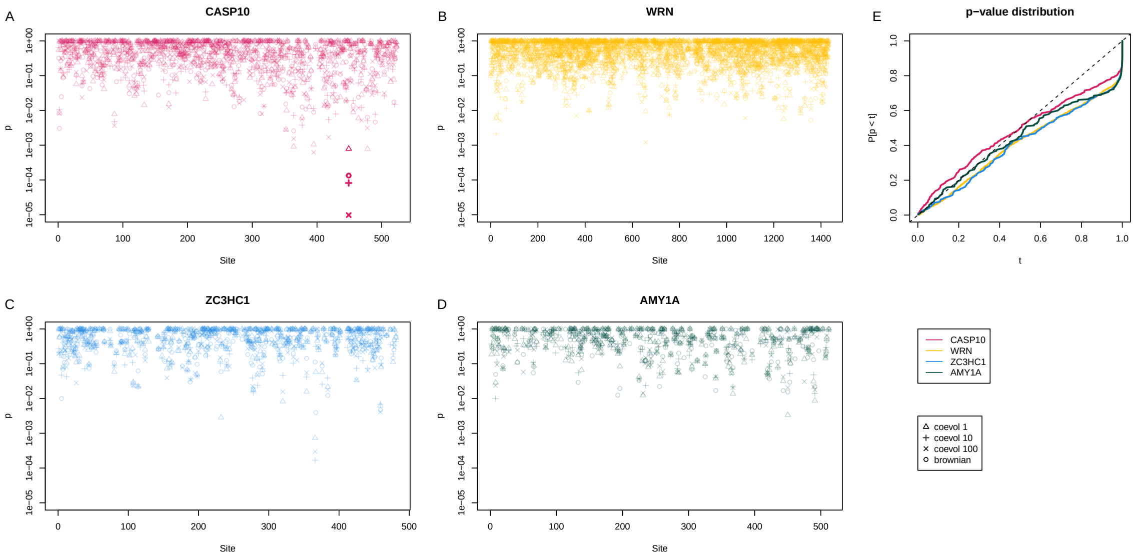


Supplementary figure 6: This figure is analogous to figure 5, but shows results obtained with model $C_S$.

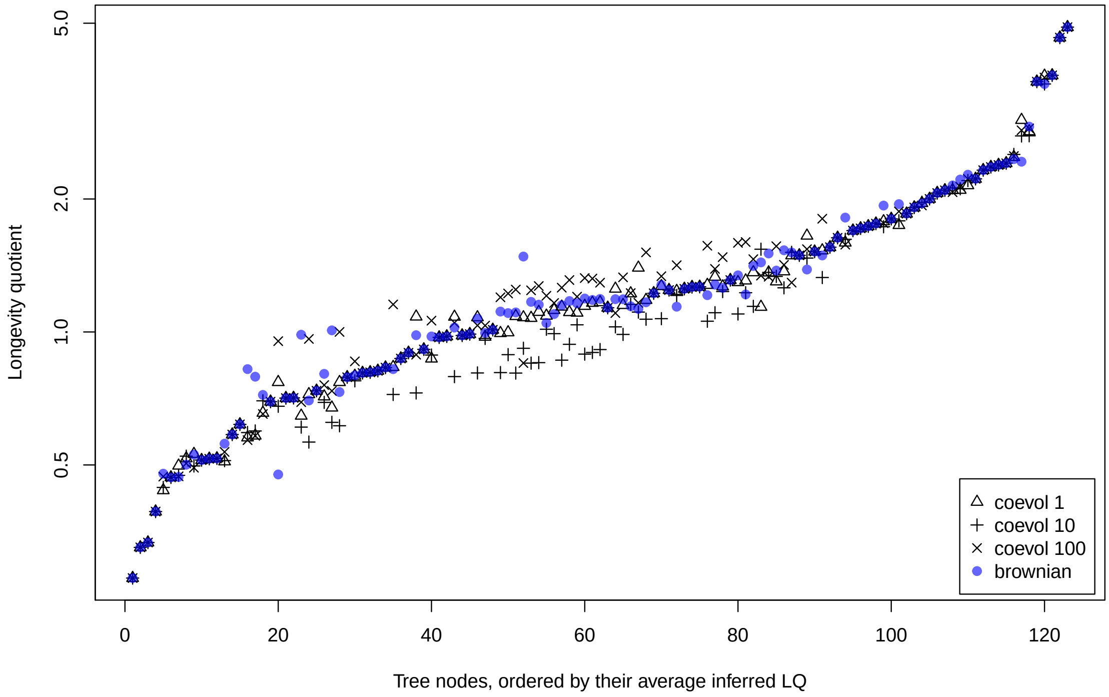


Supplementary figure 7: Inferred values of longevity quotient obtained using Coevol (and the three gene concatenates) and using Brownian motion reconstruction. Each point on the plot shows the inferred value (on the Y-axis) for a given node of the tree (on the X-axis) and given inference method (represented by the point's shape). Nodes were sorted according to the mean of the three values inferred with Coevol.

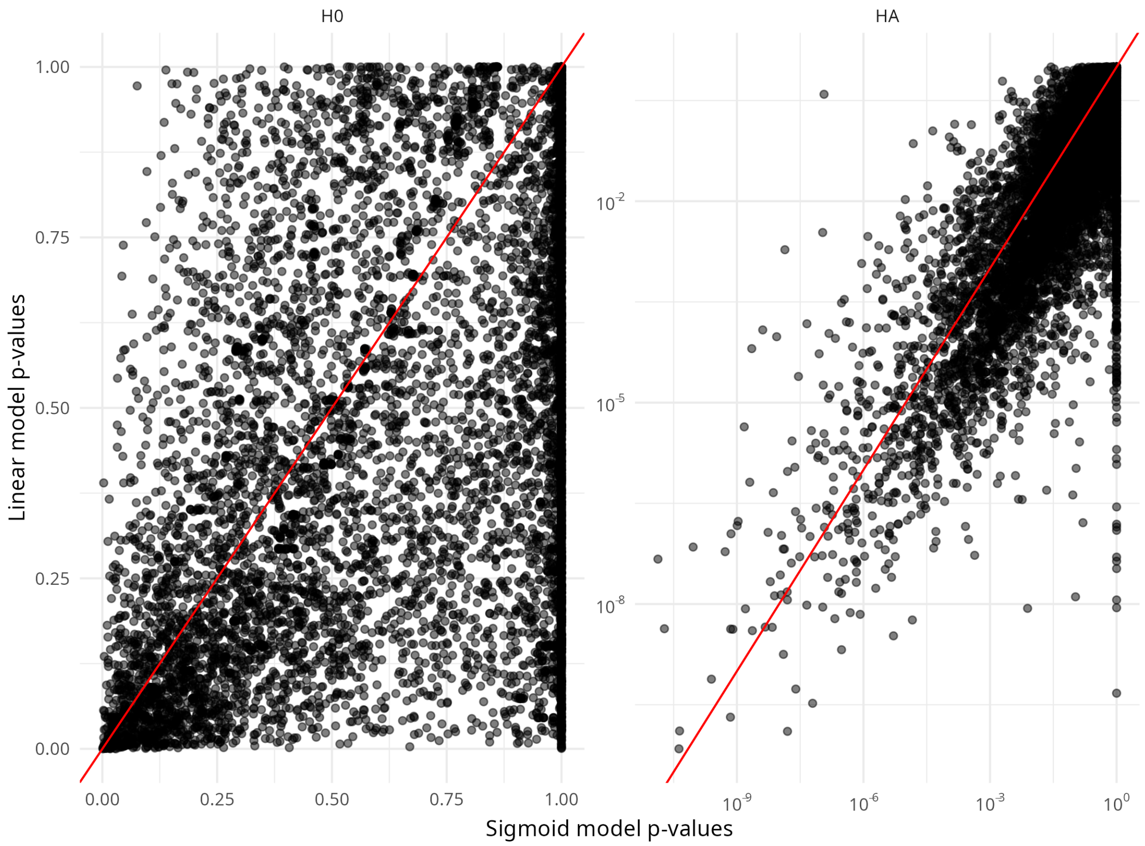


Supplementary figure 8: Comparison of $p$-values obtained with model $C_S$ and $C_L$ on simulations, under $H_0$ (left, linear scale) and $H_A$ (right, $\log_{10}$ scale).

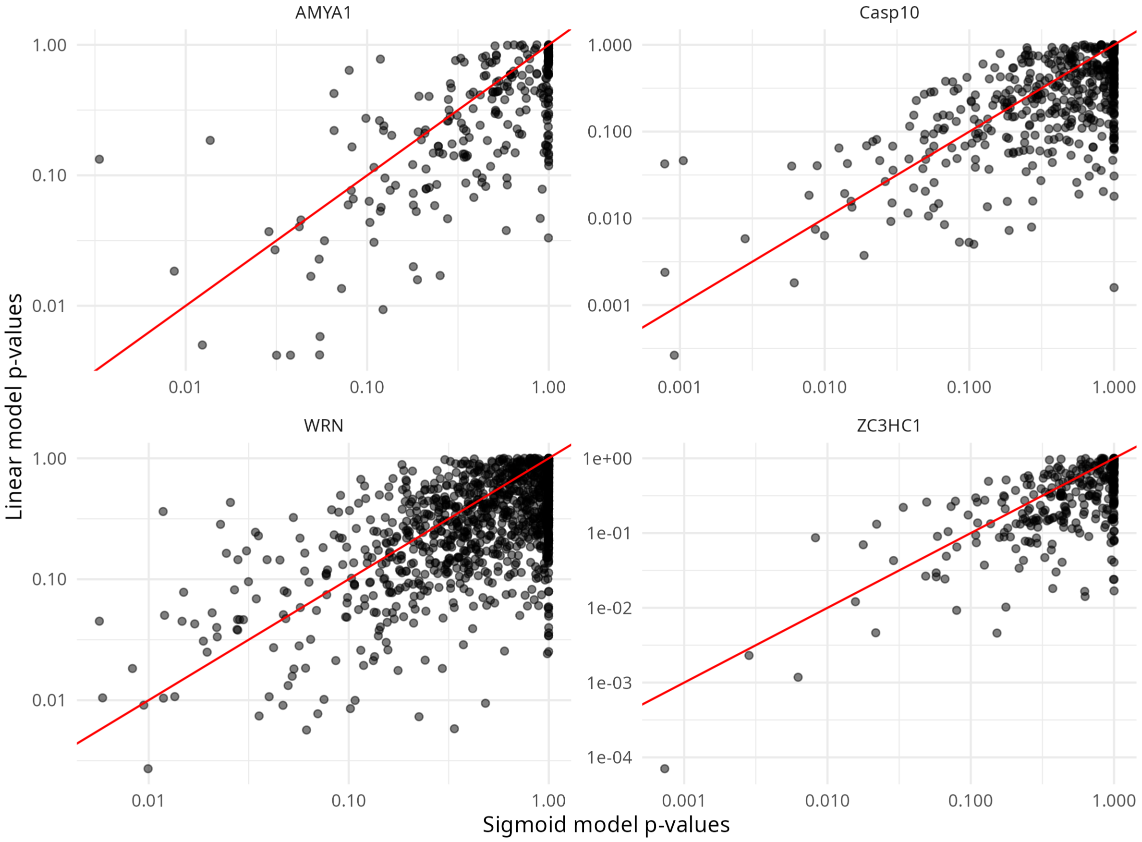


Supplementary figure 9: Comparison of $p$-values obtained with model $C_S$ and $C_L$ on empirical data.

| Species 1 | Species 2 | upper bound | lower bound |
|---|---|---|---|
| felCat8 | canFam3 | 66.09 | 37.71 |
| canFam3 | musFur1 | 66.09 | 37.71 |
| bosTau8 | panHod1 | 27.29 | 15.99 |
| panHod1 | turTru2 | 66.09 | 50.7 |
| vicPac2 | turTru2 | 66.09 | 50.7 |
| loxAfr3 | triMan1 | 65 | 54 |
| loxAfr3 | oryAfe1 | 162.5 | 56 |
| mm10 | rn6 | 15.99 | 7.25 |
| pteVam1 | myoLuc2 | 66.09 | 48.07 |
| sorAra2 | eriEur2 | 162.5 | 61.66 |
| otoGar3 | saiBol1 | 66.09 | 56 |
| ponAbe2 | hg38 | 33.9 | 11.65 |
| gorGor3 | hg38 | 33.9 | 5.33 |
| panTro4 | hg38 | 6.5 | 1.0 |
| chlSab2 | hg38 | 33.9 | 24.44 |
| monDom5 | sarHar1 | 127.2 | 48.07 |
| oryCun2 | ochPri3 | 66.09 | 48.07 |
| octDeg1 | speTri2 | 66.09 | 56 |
| oryCun2 | speTri2 | 162.5 | 56 |

Supplementary table 1: Fossil calibration used as an input to Coevol. Each line describes a constraint stating that the common ancestor of Species 1 and 2 was living in the given interval.
